# Predicting Growth and Carcass Traits in Swine Using Metagenomic Data and Machine Learning Algorithms

**DOI:** 10.1101/363309

**Authors:** Christian Maltecca, Duc Lu, Costantino Schillebeeckx, Nathan P McNulty, Clint Schwab, Caleb Schull, Francesco Tiezzi

**Affiliations:** North Carolina State University,Animal Science Department, Raleigh,27695, USA; Matatu Inc., Saint Louis, 63108, USA; The Maschhoffs LLC., Carlyle,62231, USA

## Abstract

In this paper, we evaluated the power of metagenome measures taken at three time points over the growth test period (weaning, 15 and 22 weeks) to foretell growth and carcass traits in 1039 individuals of a line of crossbred pigs. We measured prediction accuracy as the correlation between actual and predicted phenotypes in a five-fold cross-validation setting. Phenotypic traits measured included live weight measures and carcass composition obtained during the trial as well as at slaughter. We employed a null model excluding microbiome information as a baseline to assess the increase in prediction accuracy stemming from the inclusion of operational taxonomic units (OTU) as predictors. We further contrasted performance of models from the Bayesian alphabet (Bayesian Lasso) as well machine learning approaches (Random Forest and Gradient Boosting) and semi-parametric kernel models (Reproducing Kernel Hilbert space). In most cases, prediction accuracy increased significantly with the inclusion of microbiome data. Accuracy was more substantial with the inclusion of metagenomic information taken at week 15 and 22, with values ranging from approximately 0.30 for loin traits to more than 0.50 for back-fat. Conversely, microbiome composition at weaning resulted in most cases in marginal gains of prediction accuracy, suggesting that later measures might be more useful to include in predictive models. Model choice affected predictions marginally with no clear winner for any model/trait/time point. We, therefore, suggest average prediction across models as a robust strategy in fitting metagenomic information. In conclusion, microbiome composition can effectively be used as a predictor of growth and composition traits, particularly for fatness traits. The inclusion of OTU predictors could potentially be used to promote fast growth of individuals while limiting fat accumulation. Early microbiome measures might not be good predictors of growth and OTU information might be best collected at later life stages. Future research should focus on the inclusion of both microbiome as well as host genome information in predictions, as well as the interaction between the two. Furthermore, the influence of microbiome on feed efficiency as well as carcass and meat quality should be investigated.

## Introduction

The efficiency of producing saleable meat products is primarily determined by costs associated with feed and by the amount of and quality of lean meat produced^12^. Utilizing feed resources more efficiently has become a definite challenge that faces the livestock industry. Recent efforts have been devoted to identify and exploit the genomic variability of individual pigs in increasing feed efficiency^3,45^. Despite its success, this approach presents logistical as well as technical limitations related to obtaining accurate individual feed intake records^6^ as well as defining and using different feed efficiency measures^7^. Perhaps most importantly, a continued effort concentrating only on the pig variability for efficiency would inevitably lead to diminished marginal gains, incurring in concomitant losses of overall fitness and genetic diversity over time^8,9^. The amount and type of bacteria present in the gut of individuals represent a key part of all mammalian organisms^10^. The makeup of the microbiome represents a vast pool of genomic diversity that contributes to the individual physiology and health^11^. Particularly, the intestinal microbiome directly affects the degradation of carbohydrates, provides short-chain fatty acids, mitigates and alter the effect of potentially toxic compounds and produce essential vitamins^12^. The impact of environmental factors, such as nutrition^13,14^, stressors and challenges associated with weaning in pigs^15,16^ and management^17,18^, have been characterized in pigs. Nonetheless, the composition and function of a healthy microbial ecosystem have not been qualitatively and quantitatively defined and used as a tool to maximize animal health and performance^19^. Particularly, microbiome composition has yet to be studied at large scales, including large sampling conducted through several stages of the production life^20^. Within this paper, we assessed the power of metagenomic predictions based on fecal samples, to foresee growth and carcass composition in a population of healthy crossbred pigs. In doing so, we employed machinery typical of host genomic predictions, including models of the Bayesian alphabet as well as semi-parametric and machine learning algorithms.

## Results

Within this work we evaluated the effectiveness of longitudinal metagenomic data to inform prediction of growth and carcass composition in swine. For this purpose, we employed and contrasted models that have been proven successful in the genomic selection arena in order to provide the blueprint for the future routine inclusion of metagenomic information in selection programs. We evaluated the performance of the proposed models in a cross-validation setting. We further tested the overall experimental design with a mixed model based post-analysis.

### Microbiome composition overtime

The distribution of taxonomical abundance for the three time points measured (weaning, 15 weeks, and 22 weeks) in the current population has been described in detail recently by Lu and colleagues^19^. Since the objective of the current paper was not to provide the ecological landscape of the population measured, the reader is referred to that paper for more details. Briefly, at the three different stages of pig development, there were 14, 21, 29, 54, 106, and 202 identified phyla, classes, orders, families, genera, and species, respectively. For the three sampling points, 95.79–97.80% of the OTUs were classified into six phyla: *Firmicutes*, *Bacteroidetes*, *Proteobacteria*, *Fusobacteria*, *Spirochaetes*, and *Actinobacteria.* Bacteria that were in the phylum *Firmicutes* represented the majority of the total population followed by *Bacteroidetes.* To evaluate the ability of the metagenomic population to predict phenotypic measures, we conducted a preliminary analysis to investigate how different sampling times affected fecal microbiome composition. To do so, we fitted a RF model similar to the one employed for growth and carcass traits (see Methods), with the only difference that in this case the model was used to classify each observation into one of the three sampling times. We report the results of the five-fold classification in Figure 2, which depicts the normalized classification confusion matrix at weaning, 15 and 22 weeks. Individual time measurements constituted three distinct microbial populations. The accuracy of classification was in all cases extremely high (> 95%). The misclassification rate was marginally higher for 15wk and 22wk (~ 3%). This result is in line with what reported by Lu and colleagues^19^ which identified two distinct microbial enterotypes at weaning but less distinct clustering at later time points.

**Figure 1.**
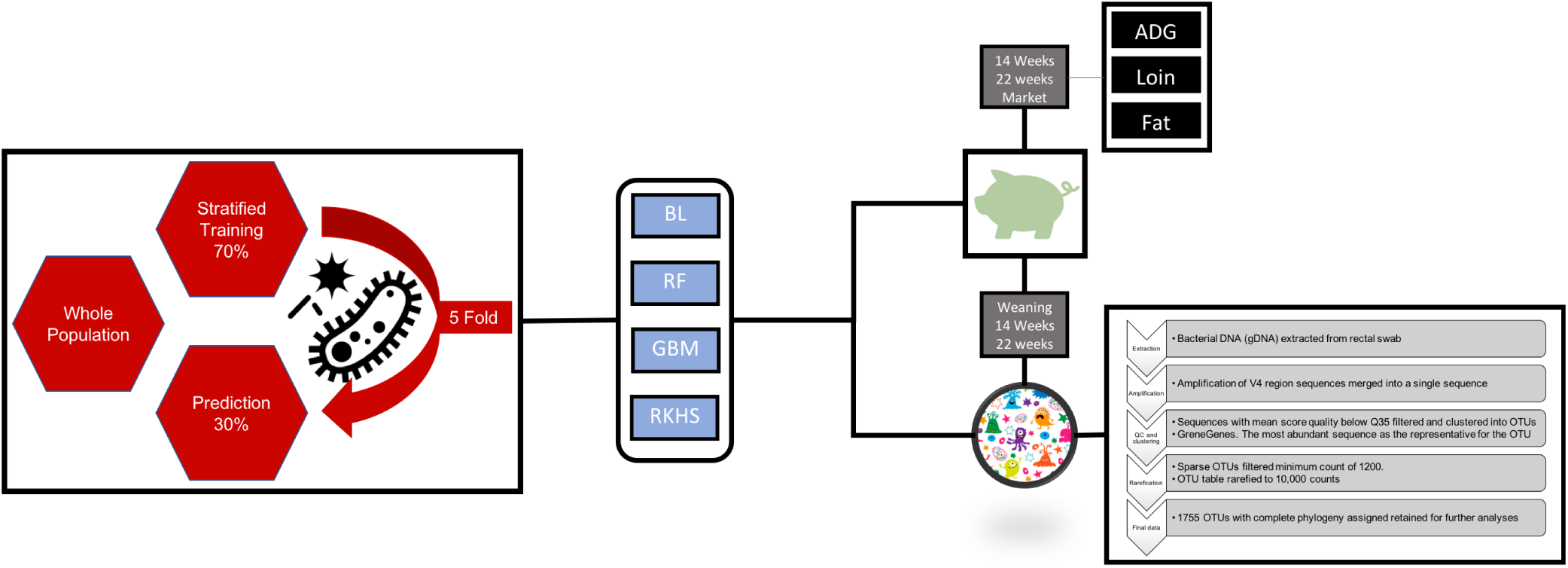
Overall Experimental design. *BL*=Bayesian Lasso, *RF*=Random Forest, *GBM*=Gradient Boosting, *RKHS*=Reproducing Kernel Hilbert Space. *ADG*=Average Daily Gain

**Figure 2.**
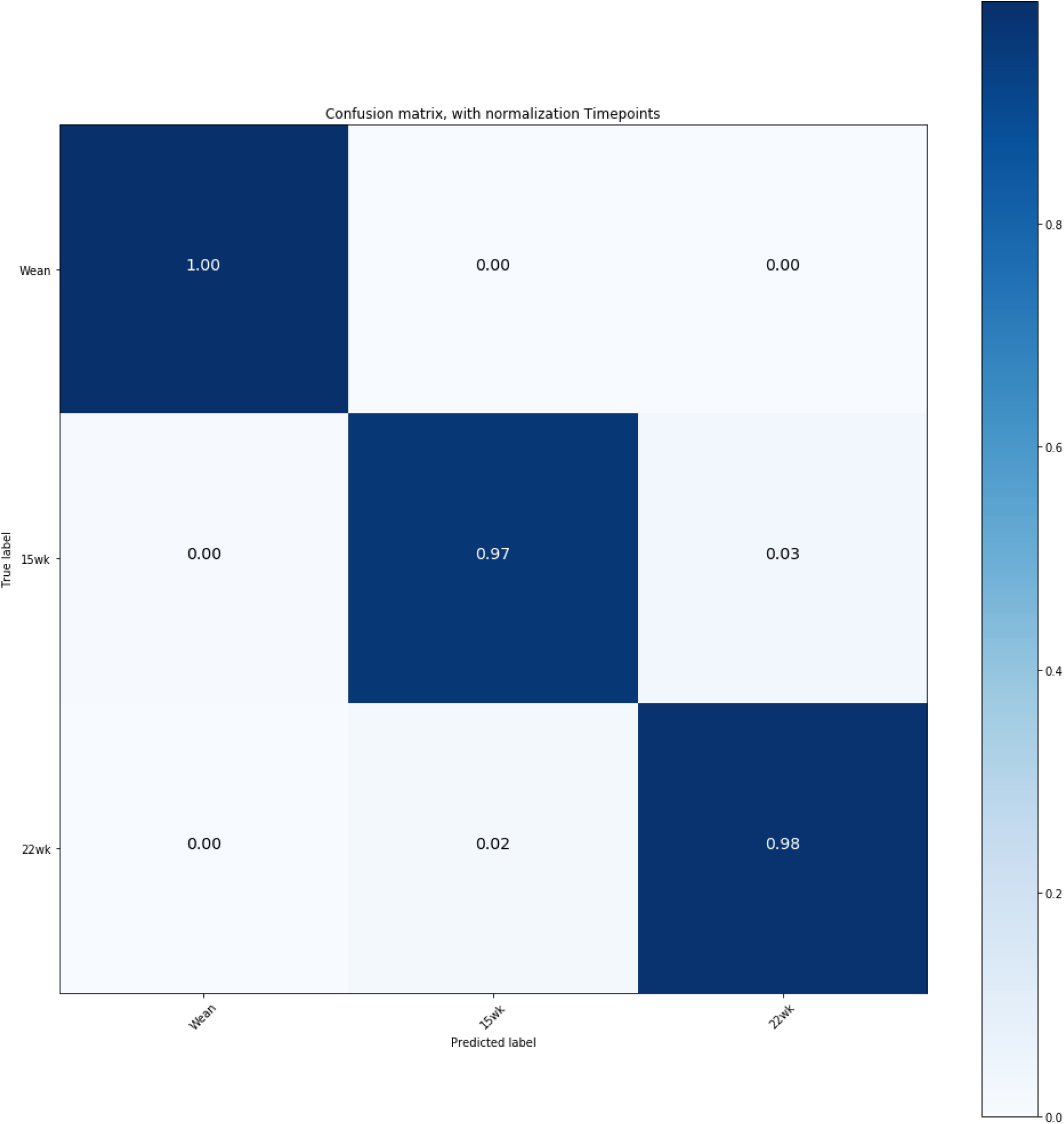
Normalized classification confusion matrix of microbiome composition at three time points. *Wean*=Weaning, *5wk*=15 weeks, *22wk*=22 weeks. Confusion matrix obtained with a RF model from a five-fold cross-validation.

### Cross-validation highlights a significant effect of microbiome for growth and carcass prediction

We first evaluated the power of metagenomic data in predicting several growth parameters in a healthy population of crossbred sires stemming from the mating of 28 founding sires’ families. For such purpose, we considered: weights, back fat, loin area and depth traits measured at 14 and 22 weeks during a growth trial; as well as daily gain measures for the same period. These were coupled with fecal microbiome information obtained for the same individuals at weaning as well as week 15 and 22 of the trial. Each trait was analyzed independently using a cross-validation scheme, in which part of the samples phenotypes and OTUs were employed to train the statistical models, and the remainder were used to validate the predictions. We considered three classes of models in the analyses: one model from the Bayesian alphabet family, Bayesian Lasso (BL)^21^; two machine learning approaches, Random Forest (RF)^22^ and Gradient Boosting (GBM)^23^; and one semi-parametric method, Reproducing Kernel Hilbert Space (RKHS)^24^. We chose the models as representative of the most widely used methods for genomic prediction in livestock and crops. We have done this to emphasize the similarity of the analyses proposed in the current work to genomic selection approaches, both in scope and methodology, as well as to provide a baseline to expand upon, with the inclusion of genomic information in future comparisons.

In Figures 3,4 and 5 are reported the accuracies of prediction for each trait, fecal microbiome time point and method combination. Microbiome contribution to prediction was measured as deviation from a null model which included only the effects of sex, sire, weight at weaning and replicate. It should be noted that the null model was fitted in all cases within each of the algorithms proposed. For ease of comparison, null models performance is nonetheless represented as the average of null models across methods. OTUs inclusion in the prediction models increased accuracies in most instances with respect to the null model. Nonetheless, the amount varied according to the microbiome time point. In general, the inclusion of microbiome composition at weaning had low predictive power for daily gain traits as well as carcass measures obtained at week 15 and 22 (Figure 3). For daily gain traits (panel A), the inclusion of metagenomic information increased accuracies of prediction of ~ 3%, yet in all cases, 90% CI of the prediction (panel C) overlapped between the null and the biom models, for all algorithms employed. Daily gain of later stages of the trial was better predicted than early growth, regardless of the inclusion of metagenomic information. Similar trends were observed for carcass traits measured at week 14 and 22, with predictions ranging from ~ 15% for loin depth (panels B, C), to ~ 40% for back fat, for both null and OTU models. Conversely, microbiome composition at week 15 increased substantially the accuracy in the test sets (Figure 4). The amount was nonetheless dependent on the trait/time combination. In general, and as expected, microbiome composition increased prediction accuracies more for traits measured concomitantly with the microbiome sampling. For daily gain traits (panel A) the inclusion of metagenomic information increased the accuracy of prediction of early growth from ~ 20 % for the null model for daily gain from birth to week 14 and from weaning to week14 to ~ 40 and 45 % for the same two traits. Similarly, for all traits measured at week 14 (panel B), microbiome information boosted prediction accuracy significantly, with gains of ~ .20 for weight and back fat and ~ .05 and .10 for loin depth and area, respectively. Similar trends were seen for week 22 traits, albeit with smaller increases and with overlapping 90 % CI (panel D) for several of the traits, with the exclusion of weight. Figure 5 depicts results of cross-validation predictions for microbiome measured at week 22. It should be noted that given the temporal succession of sampling, combinations of phenotypes measured at week 14 and microbiome at week 22 should be interpreted with caution due to the temporal succession of the measures. Again for most of the traits microbiome information increased prediction. Yet for most trait/models combinations the increase was not significant. Specifically, and focusing on week 22 traits, only weight and back fat benefitted from OTUs inclusion with gains of ~ 0.08 for back fat and ~ 0.05 for weight. Interestingly, OTU did not increase accuracy of prediction for later daily gains traits (from week 14 to week 22 and from week14 to market).

**Figure 3.**
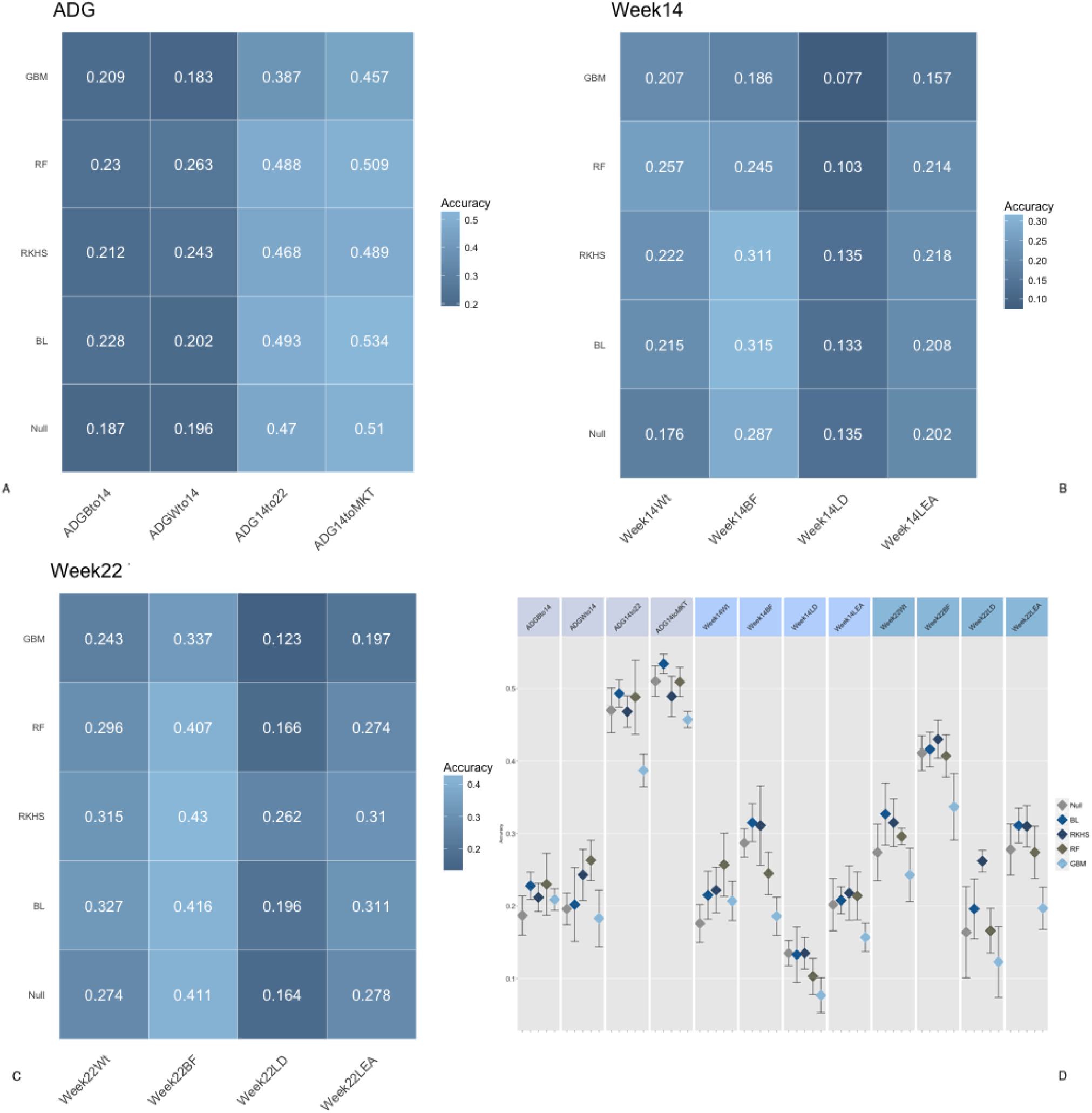
Accuracy of prediction for microbiome composition at Weaning. *Panel A*=Accuracy for daily gain traits, *Panel B*= Accuracy for Week 14 traits, *Panel C*= Accuracy for Week 22 traits, *Panel D*= 90% confidence interval for model/trait combinations. Confusion matrix obtained with a RF model from a five-fold cross-validation.*BL*=Bayesian Lasso, *RF*=Random Forest, *GBM*=Gradient Boosting, *RKHS*=Reproducing Kernel Hilbert Space. *ADGBto14*=Average Daily Gain Weaning to week14, *ADGWto14*=Average Daily Gain Weaning to week14, *ADG14to22*=Average Daily Gain week14 week22, *ADG14toMKT*=Average Daily Gain week14 to Market, *Week14Wt*=weight at week14, *Week14BF*=backfat at week14, *Week14LD=*;loin depth at week14, *Week14LEA*=loin area at week14, *Week22Wt*=weight at week22, *Week22BF*=backfat at week12, *Week22LD*=loin depth at week22, *Week22LEA*=loin area at week22

**Figure 4.**
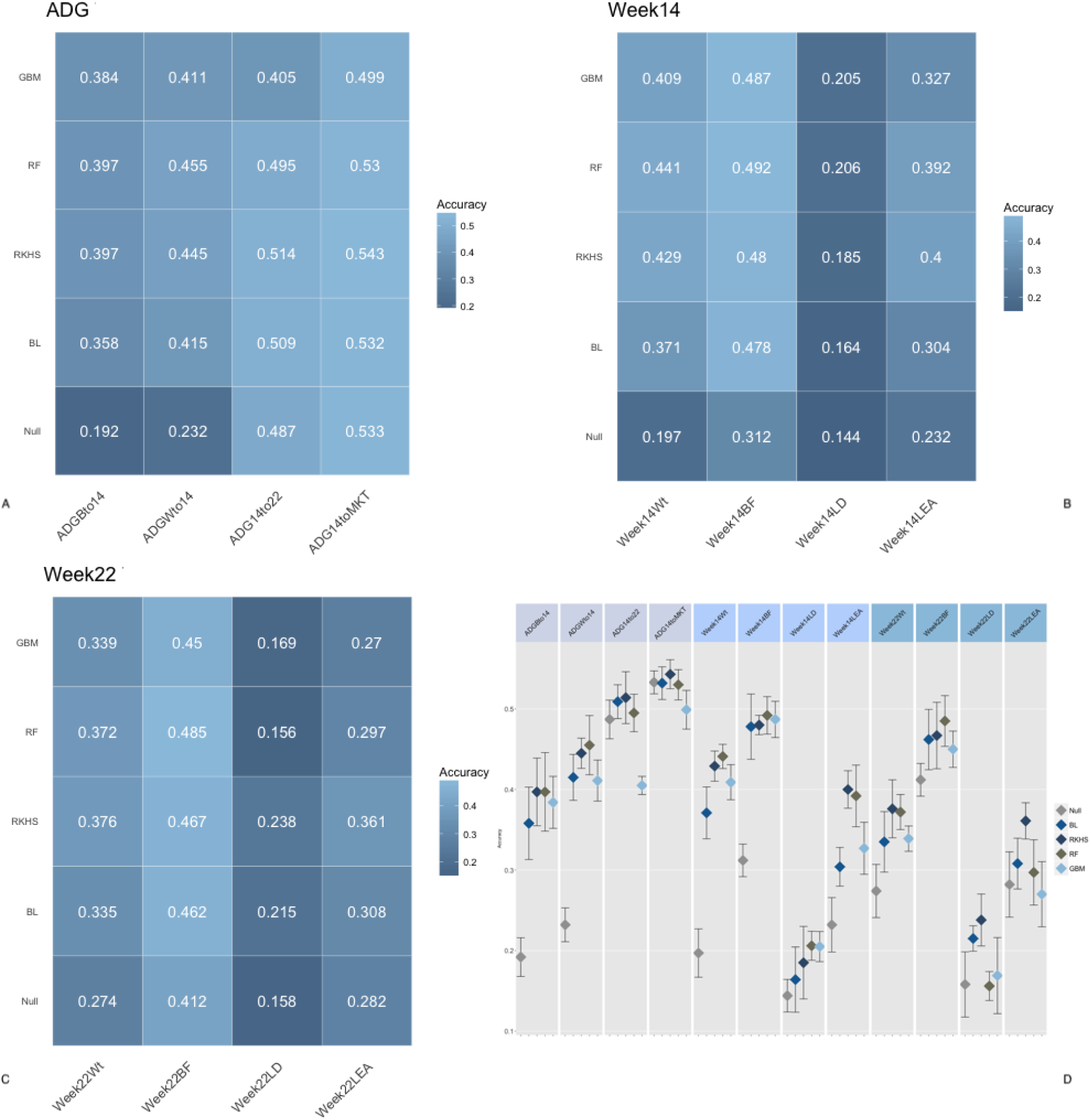
Accuracy of prediction for microbiome composition at Week 15. *Panel A*=Accuracy for daily gain traits, *Panel B*= Accuracy for Week 14 traits, *Panel C*= Accuracy for Week 22 traits, *Panel D*= 90% confidence interval for model/trait combinations. Confusion matrix obtained with a RF model from a five-fold cross-validation.*BL*=Bayesian Lasso, *RF*=Random Forest, *GBM*=Gradient Boosting, *RKHS*=Reproducing Kernel Hilbert Space. *ADGBto14*=Average Daily Gain Weaning to week14, *ADGWto14*=Average Daily Gain Weaning to week14, *ADG14to22*=Average Daily Gain week14 week22, *ADG14toMKT*=Average Daily Gain week14 to Market, *Week14Wt*=weight at week14, *Week14BF*=backfat at week14, *Week14LD*=loin depth at week14, *Week14LEA*=loin area at week14, *Week22Wt*=weight at week22, *Week22BF*=backfat at week12, *Week22LD*=loin depth at week22, *Week22LEA*=loin area at week22

**Figure 5.**
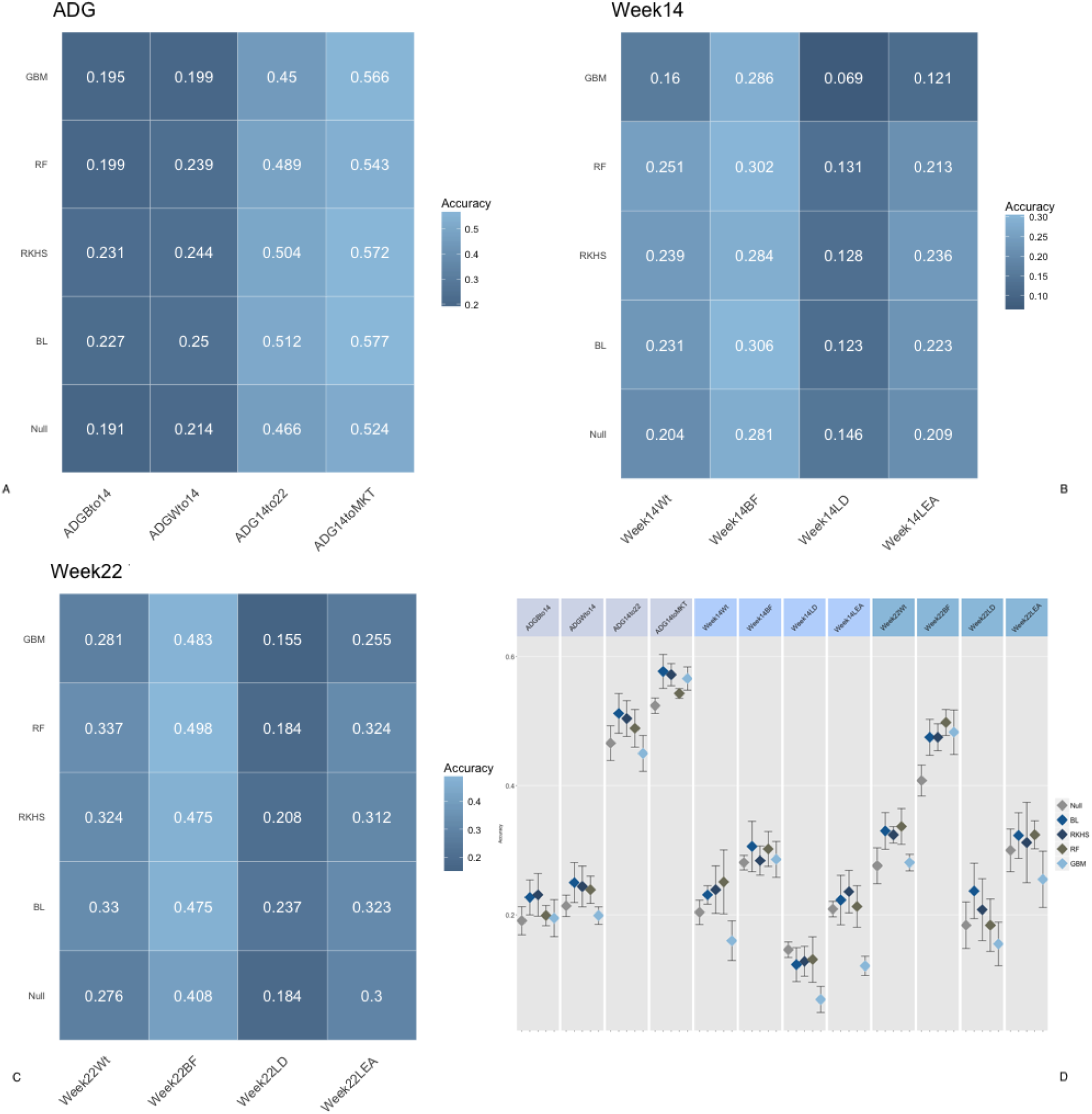
Accuracy of prediction for microbiome composition at Week 22. *Panel A*=Accuracy for daily gain traits, *Panel B*= Accuracy for Week 14 traits, *Panel C*= Accuracy for Week 22 traits, *Panel D*= 90% confidence interval for model/trait combinations. Confusion matrix obtained with a RF model from a five-fold cross-validation.*BL*=Bayesian Lasso, *RF*=Random Forest, *GBM*=Gradient Boosting, *RKHS*=Reproducing Kernel Hilbert Space. *ADGBto14*=Average Daily Gain Weaning to week14, *ADGWto14*=Average Daily Gain Weaning to week14, *ADG14to22*=Average Daily Gain week14 week22, *ADG14toMKT*=Average Daily Gain week14 to Market, *Week14Wt*=weight at week14, *Week14BF*=backfat at week14, *Week14LD*=loin depth at week14, *Week14LEA*=loin area at week14, *Week22Wt*=weight at week22, *Week22BF*=backfat at week12, *Week22LD*=loin depth at week22, *Week22LEA*=loin area at week22

The results presented are in line with what has been observed in other studies. He and colleagues^25^ found that swine gut microbiome had a moderate effect on fat with microbiome explaining from 1.5 % to 2.73% phenotypic variance for average back fat and abdominal fat weight, respectively. Similarly, Fang and colleagues^26^ found 119 OTUs associated with intramuscular fat in growing pigs. Furthermore, McCormack et al.^27^ identified several gut microbes potentially associated with porcine feed efficiency and Yang and colleagues^28^ identified two potential enterotypes in Duroc pigs associated with residual feed intake. Data on daily gain and weight is more sparse yet, for example, Ramayo et al.^29^ identified clusters of piglets significantly associated with body weight at 60d and average daily gain. It is worth noting that in most cases these studies focused on either the identification of ecological populations of bacteria or the identification of specific OTUs associated with a particular phenotype. To the best of our knowledge, this is the first attempt to rigorously characterize the overall predictive ability of the metagenome for growth and carcass traits in swine and livestock in general. In our analysis in most cases the inclusion of the microbiome composition boosted prediction accuracy beyond what expected by the identification of few important taxonomical units, not dissimilarly from what observed in genomic predictions in several livestock species^30^, suggesting a more complex interconnection between different OTUs and microbiome compositions than highlighted in previous studies. Furthermore, a growing body of literature exists pointing to a rich interplay between the pig and its metagenome^19,31^. This represents both a challenge and an opportunity to incorporate metagenomic information in selection programs effectively. The metagenome could potentially be considered an entirely environmental source of variation but also one at least partially under the direct control of the host. The methods employed in the current analysis would prove extremely flexible in integrating the full spectrum of variability generated by the availability of microbiome and host genomic data. Some of these approaches could be applied directly following GxE examples in both plants and livestock^32,33^.

### Model choice partially influence prediction accuracy with results depending on the time trait combination

We investigated the effectiveness of different model classes to incorporate metagenomic information for the prediction of growth and carcass phenotypes in pigs. We chose models ranging from completely, to semi, to non-parametric to recognize and possibly capture the complex interdependent structure of OTUs compositions. The models were tested independently for each trait time point combination. We evaluated the performance by comparing models including metagenomic composition to a baseline model including only general design factors (see Methods). Bayesian Lasso is one model of the “Bayesian Alphabet”^34^ family, that has gained popularity in genomic selection due to its ability to effectively handle large *p* small *n* problems in genomic prediction as well as providing a framework for feature selection. BL was proposed by Xu et al.^21^ and de los Campos et al.^21^. We chose it as one of the most robust and popular choices in the parametric class of models. Reproducing kernel Hilbert Space is a particularly flexible class of semi-parametric models that have been proposed to fit complex multidimensional data. They have recently gained popularity in livestock and crop breeding thanks to the work of Gianola and colleagues^35^ and of de los Campos et al.,^36^. Models of this class rely on the choice of an appropriate kernel that is then employed in models of form not dissimilar from the mixed models commonly employed in breeding settings. Random Forest is an ensemble method fitting decision trees on various sub-samples of the dataset^37^. Random forest models are generally robust to over-fitting and can capture complex interaction structures in the data^38^. Gradient Boosting is an alternative ensemble method^23^ aiming at combining predictors, in this case in a sequential manner, by forming committees of predictors with higher predictive ability than single ones. Panel D of Figures 3,4 and 5 portray point estimates and 90 % CI for each model trait combination. In the vast majority of cases the choice of model was a wash. In our analysis we weren’t able to identify a clear winner and for the most part models CI largely overlapped. Reproducing Kernel Hilbert Space models emerged as the most stable approach across scenarios in terms of ranking and magnitude of the CI,followed by Bayesian Lasso and random forest,while gradient boosting showed the largest variation in performance across trait times. At weaning gradient boosting models in some cases performed worse the null model. This is unsurprising though, as in most cases microbiome at weaning contributed little to the learning of the models. Our results are similar to what observed for the prediction of complex traits with genomic information in both plants^39^ and in livestock^40,41^, where different classes of models performed similarly over a wide variety of conditions so that in most cases the choice of model is somewhat dependent of population and data structure that the underlying biological signal. It is important to note nonetheless, that while for DNA polymorphism informed predictions, marker information is somewhat, (loosely speaking) a fixed parameter, OTU composition can be much more variable across both individuals and experimental settings, due to variability in sampling procedures, environmental conditions, as well as the bioinformatic machinery employed in obtaining taxonomical units. While we do recognize that some of this variability cannot be effectively managed through statistical modeling, we also believe that some of these models might be more flexible in handling this sources of variation. This should be the subject of further investigation, and it is beyond the scope of the current paper. Within this work and in recognition of this complexity, we attempted to overcome some of these limitations by obtaining prediction accuracies averaged across models. Results from this analysis were obtained by pooling information across replicate and methods and are presented in Figure 6. Results in this case are presented with two competing models, a null model (obtained again pooling null fit across methods) and a metagenome model (biom) obtained averaging the performance of each trait/method combination. Results for the most part recapitulate what presented in the previous section albeit in some cases, differences between null and metagenomic model have shrunk (e.g. for week 22 back fat).

**Figure 6.**
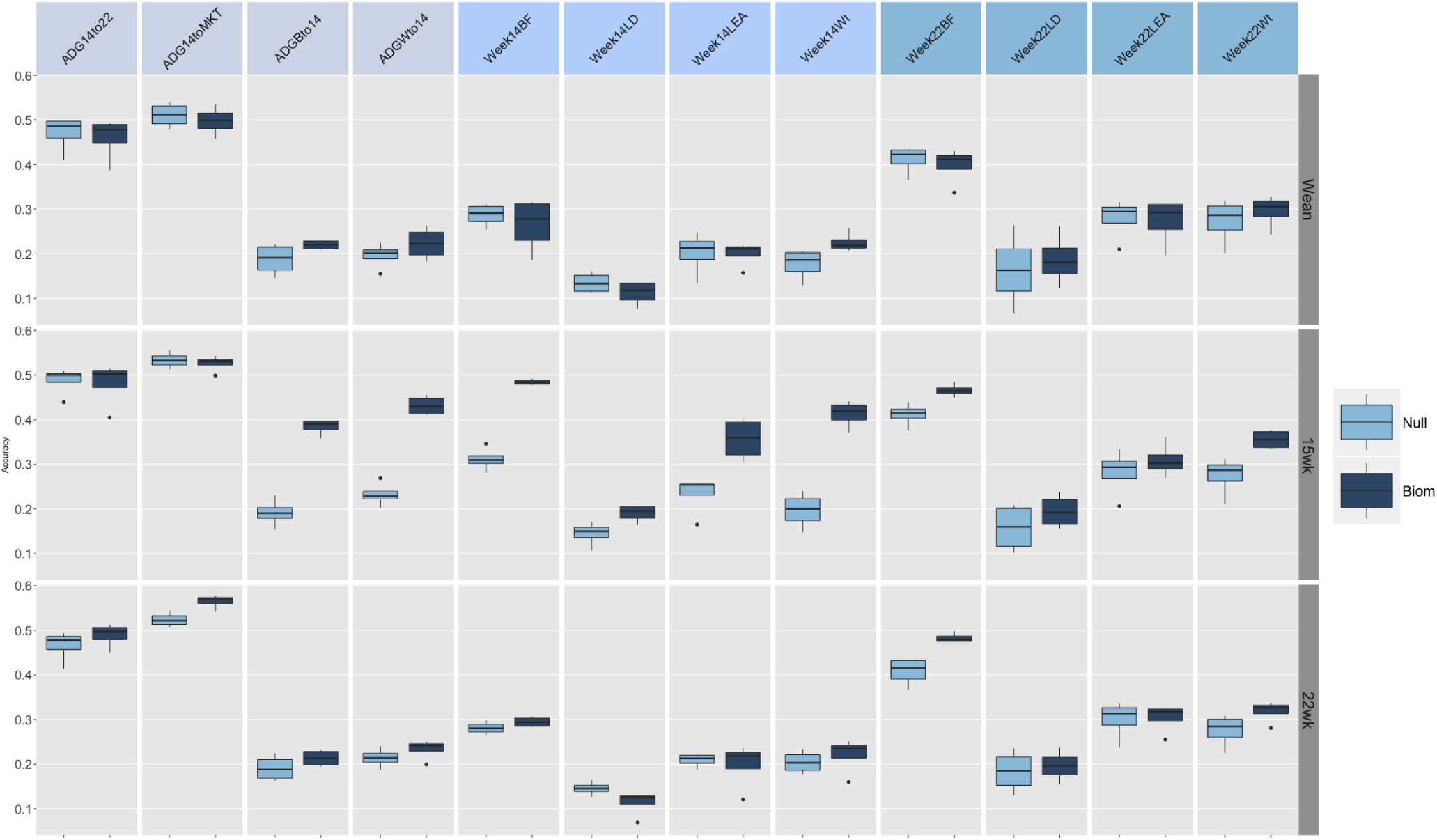
Model Average accuracy of prediction for microbiome composition at Weaning week 14 and week 22. *Null*=Average of null models. *Biom*=Average of Microbiome *modelsADGBtol4*=Average Daily Gain Weaning to week14, *ADGWtol4*=Average Daily Gain Weaning to week14, *ADG14to22*=Average Daily Gain week14 week22, *ADG14toMKT*=Average Daily Gain week14 to Market, *Week14Wt*=weight at week14, *Week14BF*=backfat at week14, *Week14LD*=loin depth at week14, *Week14LEA*=loin area at week14, *Week22Wt*=weight at week22, *Week22BF*=backfat at week12, *Week22LD*=loin depth at week22, *Week22LEA*=loin area at week22

### Post-analysis of the results

We attempted to evaluate the overall influence of all factors in the design on predictive performance with a post-analysis of the cross-validation study. To do so we employed a standard LMM approach (see Methods) and obtained least square mean estimates and contrasts for all variables in the analysis. Namely we fitted the effect of the inclusion of metagenomic information, the algorithm used for the analysis, the time point at which the fecal microbiome was sampled, the trait analyzed and all the pairwise interactions. The response variable was in this case the accuracy of prediction in the cross-validation experiment. Results of this investigation are reported in Table 2 and Figure 7. Table 2 reports the Type IIIANOVA of the overall experimental design. All factors and their interactions were highly significant with the exception of the interaction between Algorithm and Trait. The interaction between algorithm and time point was also just below the *P* < 0.05 significance threshold. Figure 7 depicts the least square means of the significant main effects and their interactions. The inclusion of metagenomic data (averaged over all other factors) increased the prediction ability of models of approximately 4 % over the null model (0.321 vs. 0.281). Of the models considered, and as seen in the previous sections GBM was the one with the lowest predictive ability (0.26) while RKHS was the one with the highest predictive power (0.32), although nearly identical to Bayesian Lasso and Random Forest algorithms. Microbiome information collected at week 15 had the highest predictive power (0.335) compared to weaning which had the lowest and week 22. Differences between the first and the two latter were ~ 5% and ~ 4%, respectively. Daily gain traits and back fat traits were the ones better predicted while loin traits, both area and depth were the ones with the lowest accuracies. The Interaction between different models and the inclusion of microbiome shows once again that RKHS models performed best regardless of the presence of metagenomic data or not. Interestingly both Random forest and gradient boosting were the algorithms that gained the most from the inclusion of OTUs information, with difference from the null model of ~ 5% in both cases. Similar trends were observed for the interaction between time point and algorithms. Finally, the interaction of metagenomic information with time points highlight how, in our data, metagenomic information collected at week 15 largely outperform (~ 10 %) all other time point (as well as null models). To the best of our knowledge, this is the first attempt to formally assess metagenomic predictions in livestock Comparable models have been used within the context of prediction of disease occurrence in human data^42^, as well as in soil microbiome associated with crop yield^43^. In both cases, the use of microbiome data achieved good predictive power, but given the vast diversity of both scope and measures, it is difficult to draw a direct comparison.

**Figure 7.**
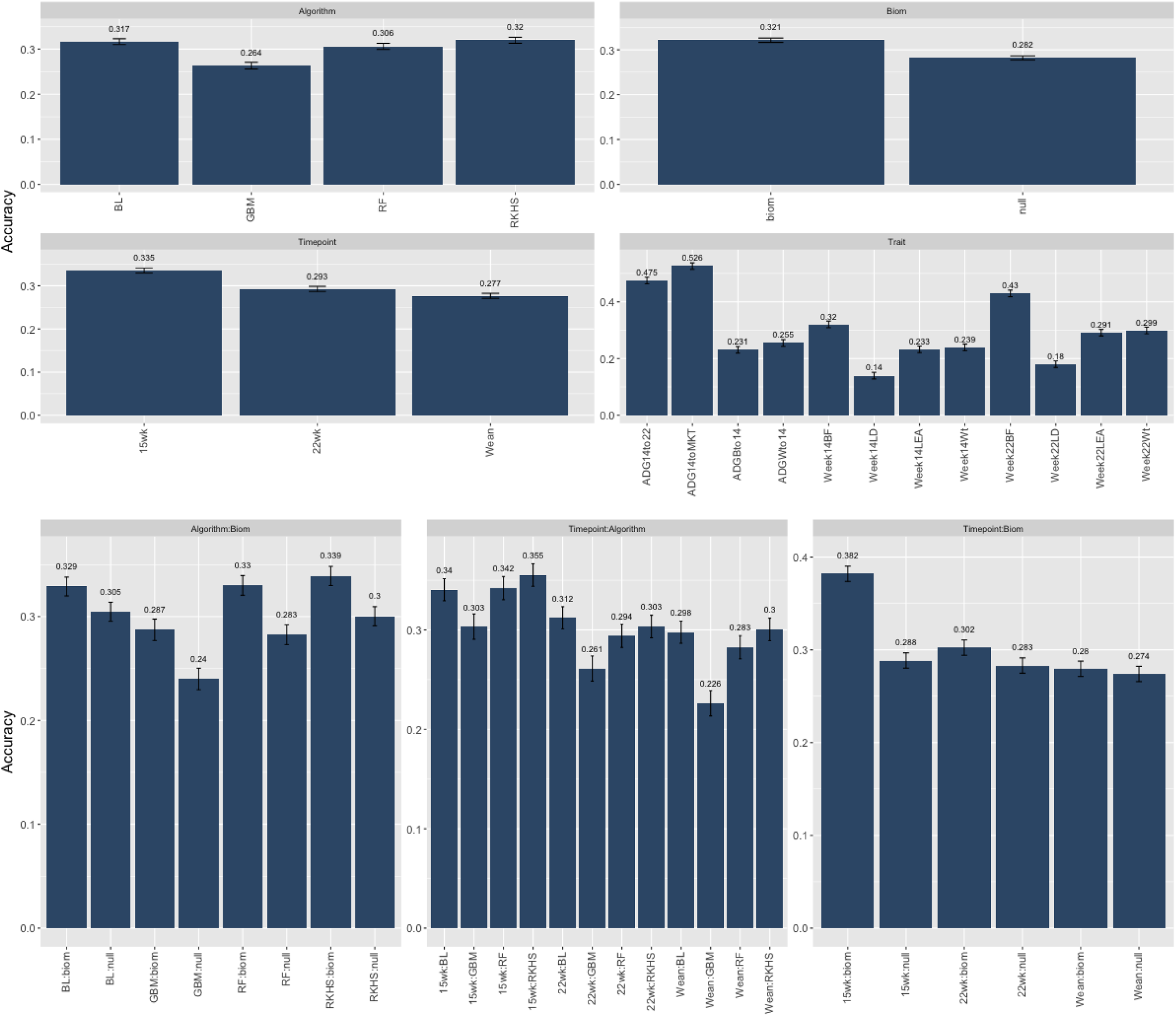
Least Square Means and SE for main effects and interactions for the post-analysis of the experimental design. *Timepoint*= 3 levels (Weaning, 15 weeks, 22 weeks), *Algorithm*=4 levels (Bayesian Lasso, Reproducing Kernel Hilbert Space, Random Forest, Gradient Boosting) *Trait*=12 levels (”ADGBto14”, “ADGWto14”, “ADG14to22”, “ADG14toMKT”, “Week14Wt”, “Week14BF”, “Week14LD”, “Week14LEA”, “Week22Wt”, “Week22BF”, “Week22LD”, “Week22LEA”), *Biom*=2 levels (null, microbiome). All elements with (:), represent pairwise interactions.

**Table 1.**
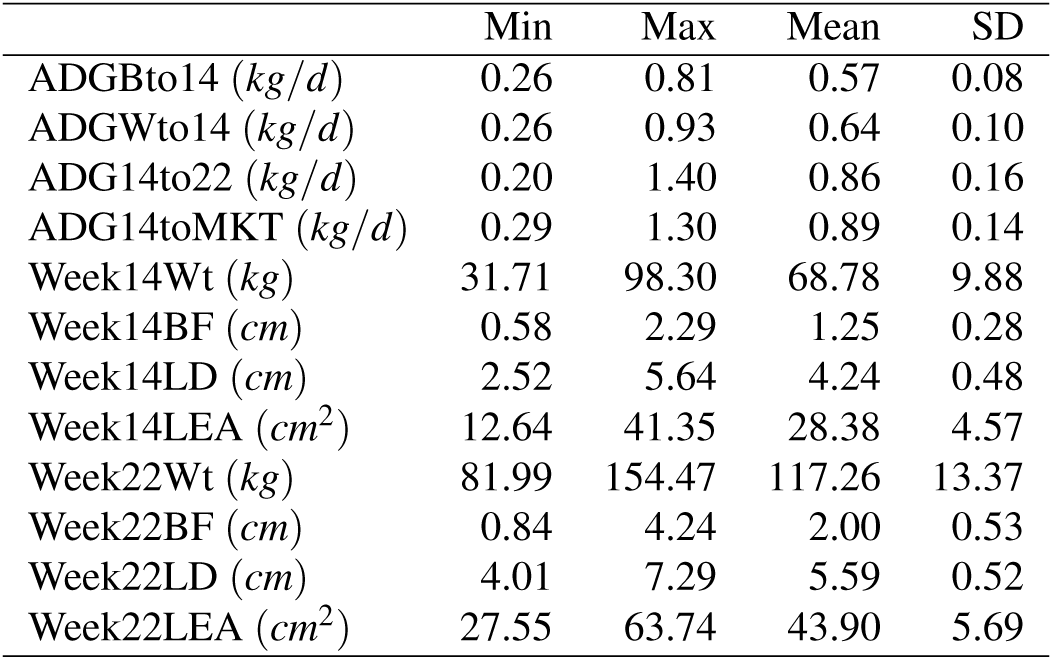
Summary of phenotypes used in the study. *ADGBto14*=Average Daily Gain Weaning to week14, *ADGWto14*=Average Daily Gain Weaning to week14, *ADG14to22*=Average Daily Gain week14 week22, *ADG14toMKT*=Average Daily Gain week14 to Market, *Week14Wt*=weight at week14, *Week14BF*=backfat at week14, *Week14LD*=loin depth at week14, *Week14LEA*=loin area at week14, *Week22Wt*=weight at week22, *Week22BF*=backfat at week12, *Week22LD*=loin depth at week22, *Week22LEA*=loin area at week22.

**Table 2.**
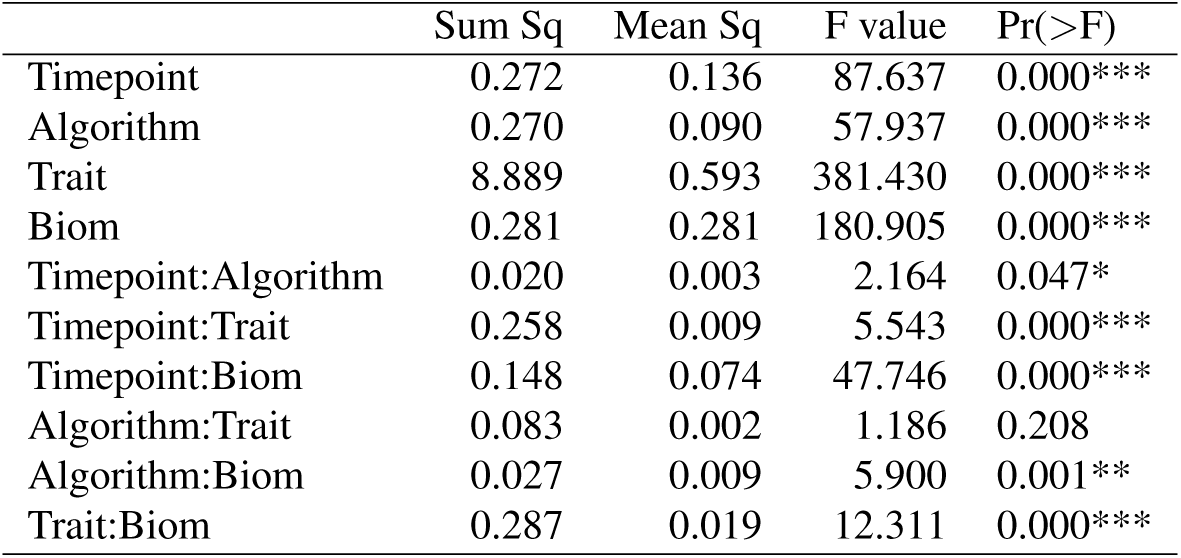
ANOVA table of the post-analysis of the experimental design. *Timepoint*= 3 levels (Weaning, 15 weeks, 22 weeks), *Algorithm*=4 levels (Bayesian Lasso, Reproducing Kernel Hilbert Space, Random Forest, Gradient Boosting) *Trait*=12 levels (”ADGBto14”, “ADGWto14”, “ADG14to22”, “ADG14toMKT”, “Week14Wt”, “Week14BF”, “Week14LD”, “Week14LEA”, “Week22Wt”, “Week22BF”, “Week22LD”, “Week22LEA”), *Biom*=2 levels (null, microbiome). All rows with (:), represent pairwise interactions.

## Discussion

In general, our cross-validation highlighted good predictive power, however results varied considerably between metagenome time sampling as well as trait measured. From our study it appears that sampling time might be a crucial factor in integrating metagenomic information in predictive models for growth. Our data suggest that samples measured in the middle of the growth trial would provide the highest amount of information. Conversely early measures of microbiome composition might not be as informative. This is somewhat in contrast with some of recent studies^28,29^ that have found different enterotypes related growth traits at earlier stages. In our experience and how highlighted by Lu et al.^19^, clustering of individuals at early time points could be the results of piglet adjusting more or less rapidly to the change in diet that normally happens at weaning. We believe this should be investigated further. Within this paper we considered the study of each time point as separate and independent. This is a simplification that allowed to build an easy cross-validation experiment to test different variables. Nonetheless, the use of longitudinal models in the future would provide a much more powerful way to investigate the importance in changes in microbiome composition, and their implication in efficient growth in livestock. To this point, some of the deep learning models developed in the context of prediction of longitudinal data^44^ should allow for a much better understanding of the complex interplay between changes in OTUs composition and the prediction of phenotypes. Nonetheless, a much larger number of individuals as well as deeper sampling would be needed to reach the necessary data granularity to make these approaches appealing. In our studies both growth traits and fatness traits achieved good predictive power. To further explore the role metagenomic information in efficiency of growth data on feeding (both amount and behavior) should be investigated. Furthermore, the current study was conducted within a single crossbred population. For the effective exploitation of metagenomic variability in pigs a larger number of populations/breeds should be investigated, given the large variability in OTU composition existent within the swine specie^45^. Within this work we have established a framework that could later be expanded to include not only metagenomic information but also host genomic data^46^, to better characterize and possibly manage the environment as well as to account for the complex relationships between host and guest variability. Microbiome composition can be effectively used as a predictor of growth and composition traits, particularly for fatness traits. Inclusion of OTU predictors could potentially be used to promote fast growth of individuals while limiting fat accumulation. Early microbiome measures might not be good predictors of growth and OTU information might be best collected at later life stages.

## Animals

The pigs used in this study were grown in a commercial setting operated by The Maschhoffs LLC (Carlyle, IL, USA). Animal use approval was therefore not needed for the data collection. Offspring for the current study originated from twenty-eight purebred Duroc sires, from a Duroc population under selection for lean growth, mated to Large White x Landrace or Landrace x Large White sows. The resulting offspring were weaned at 18.6 days (±1.09) and subsequently moved to a nursery-finishing facility. Here individuals were grouped in batches of 20 pigs per pen. Pen mates were paternal half-siblings of the same gender and similar weaning weight. We performed six replicates of this basic experimental block, each composed of 2 pens (one pen of female and one pen of castrated male pigs) from each of the 28 sires. The test period began the day the pigs entered to the nursery-finishing facility. Individuals were fed a standard pellet diet during nursery, growth, and finish periods they were fed standard pelleted feed while during the grow-finish period, a standard diet based on sex and live weight was fed. Diet formulations and their nutritional values are provided [see Additional file 1]. The pigs received a standard vaccination and medication routine [see Additional file 2]. End of test was reached on a pen-specific basis when all pigs in a pen achieved an average live weight of 136 kg. Their average age at harvest was 196.4 days (±7.86). We collected rectal swabs from all pigs in a pen at three time points: weaning, 15 weeks post weaning (average 118.2 ±1.18 days, hereafter “wk15”), and 22 weeks post weaning (average 196.4 days ±7.86 days, hereafter “wk22”). Four pigs were chosen randomly per pen for lean carcass growth measurements, and their rectal swabs were used for microbiome sequencing. In the end, the number of samples at weaning, week 15, and week 22 were 1205, 1295, and 1283, respectively. There were 1039 animals with samples collected at all 3 time points. More details on the distribution of samples across families, time points, and sex are provided [see Additional file 3]. Loin depth, loin area as well as back fat thickness and weights were recorded on live animals at weeks 14 and 22 post-weaning and at market weight. These measures will be hereafter referred to as Week14LEA, Week14LD, Week14BF Week14Wt and Week22LEA, Week22LD, Week22BF, Week22Wt, respectively. Likewise, average Daily gain were measured as difference in live weight from birth to week 14 (ADGB14), from weaning to week 14 (ADGW14) from week 14 to week 22 (ADG1422) and from week 14 to market (ADG14MKT). A summary of the traits employed in the current analysis is reported in Table 1.

## DNA extraction and purification

Total DNA (gDNA) was extracted from each rectal swab by mechanical disruption in phenol:chloroform. Briefly, *μL* 650 of extraction buffer (200 mM Tris; 200 mM NaCl; 20 mM EDTA, pH 8.0) was added to each swab stored in a 2 mL self-standing screw cap tube (Axygen, CA, USA). Tubes were shaken using a Mini-BeadBeater-96 (MBB-96; BioSpec, OK, USA) for 20 s to free sample material from the swab head. Following a brief centrifugation (10 s; 500 x g) to pull down any dislodged material, each swab head was removed from its tube using sterile forceps. Samples were frozen solid at –80°C, and approximately 250 *μL* of 0.1 mm zirconia/silica beads (BioSpec) and a 3.97 mm stainless steel ball were added to the sample (while still frozen, to avoid splashing). Samples were allowed to thaw briefly, after which 210 *μL* 20% SDS and 500 *μL* phenol:chloroform:IAA (25:24:1, pH 8.0) were added. Bead-beating was performed on the MBB-96 (4 min; room temperature), samples were centrifuged (3,220 x g; 4 min), and 250 *μL* of the aqueous phase was transferred to a new tube. 100 *μL* of this crude DNA was then further purified using a QIAquick 96 PCR purification kit (Qiagen, MD, USA). Purification was performed per the manufacturer’s instructions with the following minor modifications: (i) sodium acetate (3 M, pH 5.5) was added to Buffer PM to a final concentration of 185 mM to ensure optimal binding of genomic DNA to the silica membrane; (ii) crude DNA was combined with 4 volumes of Buffer PM (rather than 3 volumes); and, (iii) DNA was eluted in 100 *μL* Buffer EB (rather than 80 *μL*).

## Illumina library preparation and sequencing

Phased, bi-directional amplification of the V4 region (515–806) of the 16S rRNA gene was employed to generate indexed libraries for Illumina sequencing using the strategy described in^47^. Amplicon libraries were quantified using the Qubit dsDNA assay kit (Thermo Fisher Scientific Inc., MA, USA) before being pooled in equimolar ratios. These final pools were purified using Agencourt AMPure XP beads (Beckman Coulter) per the manufacturer’s instructions. Purified pools were supplemented with 5–10% PhiX control DNA and were sequenced on an Illumina MiSeq machine as paired-end 2×250 + 13bp index reactions using the 600v3 kit. Un-demultiplexed FASTQ files were generated by MiSeq Reporter. All sequencing was performed at the DNA Sequencing Innovation Lab at the Center for Genome Sciences and Systems Biology at Washington University in St. Louis.

## 16S rRNA gene sequencing and quality control of data

Pairs of V4 16S rRNA gene sequences were first merged into a single sequence using FLASh v1.2.11^48^, with a required overlap of at least 100 and not more than 250 base pairs in order to provide a confident overlap. Sequences with a mean quality score below Q35 were then filtered out using PRINSEQ v0.20.4^49^. Sequences were oriented in the forward direction and any primer sequences were matched and trimmed off; during primer matching, up to 1 mismatch was allowed. Sequences were subsequently de-multiplexed using QIIME v1.9^50^. Sequences with > 97% nucleotide sequence identity were then clustered into operational taxonomic units (hereafter “OTUs”) using QIIME with the following settings: `max_accepts = 50`, `max_rejects = 8`, `percent_subsample = 0.1` and --`suppress_step4`. A modified version of GreenGenes (The Greengenes Database Consortium^51,52,53^) was used as the reference database. Input sequences that had 10% of the reads with no hit to the reference database were then clustered de novo with UCLUST^54^ to generate new reference OTUs to which the remaining 90% of reads were assigned. The most abundant sequence in each cluster was used as the representative sequence for the OTU. Sparse OTUs were then filtered out by requiring a minimum total observation count of 1200 for an OTU to be retained, and the OTU table was rarefied to 10,000 counts per sample. Average Good’s coverage estimates for samples at weaning, week 15 and week 22 were 0.99 ± 0.002, 0.98 ± 0.002, and 0.98 ± 0.002, respectively. Finally, the Ribosomal Database Project (RDP) classifier (v2.4) was retrained in the manner described in Ridaura and colleagues^55^ with 0.8 cutoff used to assign taxonomy to the representative sequences. After data processing and quality control 1755 OTUs were available for further analyses.

## Statistical analysis

### Training and testing sets

A stratified five-fold cross validation scheme was used to recursively randomly split data into training (~ 70% of observations) and prediction (~ 30% of observations) sets, maintaining equal representation of the 28 sires present in the trial. For the investigation, each combination of method, trait and time was treated as a separate analysis and accuracy of prediction was obtained as the average correlation between predicted and measured phenotypes in the test sets. A pictorial representation of the overall experimental design is depicted in Figure 1.

### Models

#### Bayesian Lasso: For each fold/trait/time point combination two models were fitted

A null model (*null*):

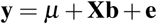

where: **y** was one of traits mentioned in the previous section,***μ*** was a population mean, **b** was a vector of fixed effects which included: *sex* (2 levels), *replicate* (6 levels), *sire* (28 levels), plus the covariate of weight at weaning, **e**, was a vector of random residuals assumed 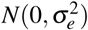 and **X** was an incidence matrix relating observations to fixed effects.

A model including the microbiome (*biom*):

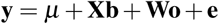

where: **o** was a vector of OTUs effects (1755 levels), **W** was a matrix of centered and scaled OTUs counts and the remainings were as in the previous model.

We fitted the BL regression model as implemented by the R^56^ package `BGLR`^57^. OTU counts were fitted to the model with the use of a double exponential prior distribution. BGLR models double-exponential density as a mixture of scaled normal densities. In the first level of the hierarchy, marker effects are assigned independent normal densities with null mean and OTU-specific variance parameter 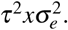 The residual variance was assigned a scaled-inverse Chi-square prior density. BGLR provides a convenient way to choose priors shape through the `R2` flag. R2 can roughly be interpreted as the expected variance proportion explained by the effect included in the model. For the residual effects default degrees of freedom of 5 were employed and an R2 of .60. Prior scale parameter where then obtained as *Sp* = *Var*(*y*)(1 – *R*2)(*d f p* + 2), with *Sp* and *d f p* the scale and degrees of freedom, respectively. OTUs specific scale parameters, τ^2^ are assigned *IID* exponential densities with rate parameter *λ*^2^/2. The hyper parameter *λ* was in this case fixed and its value was assigned through a grid search on the full dataset/trait combinations (results not shown).

#### Random Forest

The general form of the null model employed here was (following González-Recio and Forni^38^):

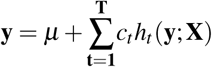

while the biom model was:

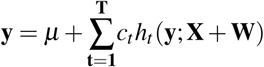

Each tree *h_t_* (*y*; *X*) or *h_t_* (*y*; *X* + *W*) for *t* ∈(1, *T*) was constructed from a random sample of the original data, and at each node a subset of features were randomly selected to create the splitting rule. Each tree was grown to the largest extent possible until all terminal nodes were maximally homogeneous^38^. The parameter *c_t_* is a shrinkage factor averaging the trees. The quality of split in RF can be measured through different criteria. For the current analysis mean square error (MSE) was employed. The remaining parameters of RF models in this work were set as follows: i) the number of trees was set equal to 1500; ii) the number of features to consider when looking for the best split was equal to the root of the number of original features. The `bigrf` package^58^ of R^56^ was used to fit RF models to the data.

#### Gradient Boosting

The general form of the null model employed here was (again following Gonzalez-Recio and Forni^38^):

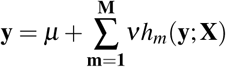

while the biom model was:

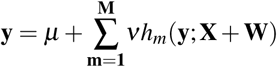

Each predictor *h_m_*(*y*;*X*) or *h_m_*(*y*;*X* + *W*) for *t* ∈(1,*M*) was, in this case, applied consecutively to the residual from the committee formed by the previous ones, the bagging step remaining similar to what described before. The `gbm` package^59^ of R^56^ was used to fit GBM models to the data. A gaussian loss function was employed. Other parameters in the GBM models were set as follow: i) the number of trees was set equal to 1500; ii) the interaction depth was set at 3; iii) the shrinkage parameter *ν* was set at 0.01.

#### Reproducing Kernel Hilbert Space

Two RKHS models were fitted:

A null model (*null*):

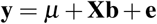

and a (*biom*) model of form:

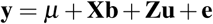

where **Z** is an incidence diagonal matrix of order(1039×1039) and **u** is a random vector of pig effects assumed 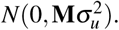 **M** was the kernel matrix based on microbiome composition, and its computation was as follows: microbiome was used at the OTU level to compute the Jensen-Shannon distance between pairs of samples, 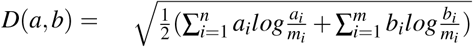 in which *D*(*a*, *b*) was the distance between samples *a* and *b*; *n* was the number of OTUs (n=1755); *a_i_* and *b_i_* were the counts of *OTU_i_* in samples *a* and *b*, respectively; *m_i_* = (*a_i_* + *b_i_*)/2^60^. The resulting square matrix (hereafter “JSD”) had zero on the diagonal, and values ranging between 0 and 1 on th off-diagonal. The **M** matrix was obtained as 1 – JSD.The RKHS regression model was implemented with the R package `BGLR` within a bayesian setting. Prior for 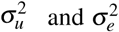 where chosen as highlighted in the previous section. R2 values for the the two parameters were set at 0.3 and 0.6, respectively.

### Post-analysis

In order to provide a comprehensive assessment of all the factors in the design we conducted a post-analysis of the experiment with the use of a standard Linear Mixed Model (LMM). All combinations of replicate/trait/method were pooled in a single dataset. The following LMM was then fitted

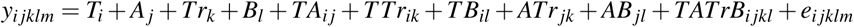

where *y_ijklm_* is the accuracy of each replicate/trait/method combination; *T_i_* is the fixed effect of the microbiome timepoint measurement (3 levels: wean, 15wk, 22wk); *A_j_* is the fixed effect of the algorithm used (4 levels: BL, RKHS, RF, GBM); *Tr_k_* is the fixed effect of the trait (12 levels: ADGBto14, ADGWto14, ADG14to22, ADG14toMKT, Week14Wt, Week14BF, Week14LD, Week14LEA, Week22Wt, Week22BF, Week22LD, Week22LEA); *B_l_* is the fixed effect of the microbiome inclusion (2 levels: null, biom); *TA_ij_ TTr_ik_ TB_il_ ATr_jk_* and *AB_jl_* are the pairwise interactions of the main effects; *TATrB_ijkl_* is the random interaction effect of *T*, *A*, *Tr* and *B* assumed 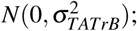 and *e_ijklm_* is the random residual effects assumed *N*(0, *σ*^2^). The LMM model was fitted with the R^56^ package `lme4`^61^. Type IIIANOVA table, least square means and contrasts were obtained with the R package `lmerTest`^62^.

## Acknowledgements (not compulsory)

We would like to thank Jessica Hoisington-Lopez from the DNA sequencing Innovation Lab at the Center for Genome Sciences and Systems Biology at Washington University in St. Louis for her sequencing expertise and Nicholas S. Grohmann for phenotype and sample collection.

## Availability of data and materials

The datasets generated and/or analyzed during the current study are not publicly available due to third-party ownership but are available from the corresponding authors upon reasonable request. All scripts used for the analysis and manuscript preparation are available from the corresponding authors upon reasonable request.

## Authors’ contributions

CM designed and carried out the analyses and drafted the manuscript. CS and NM were responsible for sequencing and bioinformatic work, as well as drafting related sections in the “Methods” section. FT and DL were involved in designing the experiment and providing consultation for the analyses. All co-authors provided comments for the manuscript. CM directed the overall research project. All authors have read and approved the final manuscript.

## Ethics approval

Phenotypic records presented in this study came from field data. Procedures for fecal sample collection adhered to the guidelines of Institutional Animal Care and Use Committee, North Carolina State University, and National Pork Board.

## Consent for publication

Not applicable.

